# Moremi Bio Agent: Application of A Foundation Model and End-to-End Automation in the Design and Validation of Monoclonal Antibodies Targeting *Plasmodium falciparum* Invasion Complex

**DOI:** 10.1101/2025.02.12.637967

**Authors:** Darlington Akogo, Jeremiah Ayensu, Nana Sam, Gertrude Hattoh, Prince Nyarko, Solomon Eshun, Mohammed Alhasan, Henrietta Mensah-Brown, Peter Quashie

## Abstract

Malaria remains a significant global health challenge, with *Plasmodium falciparum* responsible for the majority of severe cases and fatalities. Targeting the parasite’s invasion mechanisms offers a promising therapeutic strategy. In this study, we leveraged a novel agentic foundational model, Moremi Bio Agent, to design monoclonal antibodies targeting the AMA1-RON2 complex, a critical component in the parasite’s invasion of human red blood cells. Using advanced structural modeling, we generated 999 antibodies, which were evaluated for binding affinity, structural integrity, and physicochemical properties. Binding affinity analysis using PRODIGY identified 864 antibodies with successful target interactions, exhibiting binding free energies (ΔG) ranging from -116.8 kcal/mol to -5.6 kcal/mol. The strongest candidates demonstrated exceptionally tight binding, with dissociation constants (K_d_) in the femtomolar to attomolar range, indicative of highly stable interactions. Additionally, structural validation confirmed that the antibodies were thermodynamically stable with robust fold reliability, essential for functional efficacy. Epitope mapping revealed highly conserved regions within the target complex, enhancing the likelihood of cross-strain efficacy. Glycosylation analysis identified key sites that could improve antibody stability and immune recognition, while BLAST comparison with known therapeutic antibodies demonstrated significant homology, underscoring their potential for clinical development. This study highlights the power of generative AI-driven computational pipelines in antibody discovery, providing a scalable and cost-effective framework for therapeutic development. The findings establish a foundation for experimental validation and optimization, with the potential to advance novel interventions against malaria and other infectious diseases.

## 1. Introduction

Malaria remains one of the most significant global health challenges, causing an estimated 608,000 deaths in 2022, with most of these deaths occurring in the WHO African Region [1]. The disease is caused by protozoan parasites of the genus Plasmodium, with Plasmodium falciparum being the most lethal species [2]. Other species that are known to cause malaria in humans are Plasmodium vivax, Plasmodium ovale, Plasmodium malariae, and Plasmodium knowlesi.

Several efforts to improve public health have led to the development of antimalarial drugs and vaccines such as Artemisinin-based combination therapies, and Mosquirix [3,4]. Nevertheless, the emergence of drug resistance, limited vaccine efficacy, and challenges in ensuring widespread access to these interventions have prompted the need for new therapeutic approaches to effectively fight against malaria [5,6]. Neutralizing the parasite’s mechanisms of red blood cell invasion represents a promising strategy for therapeutic intervention.

One key protein complex which plays an essential role in the invasion process of P. falciparum is AMA1-RON2. The AMA1-RON2 complex facilitates the formation of a tight junction between the parasite and the host red blood cell, a critical step for invasion [7]. This study is critical as would other critical complexes including RH5-CyRPA-Ripr [8]. These complexes are highly conserved and indispensable, making them attractive targets for neutralizing monoclonal antibodies (mAbs) aimed at blocking invasion and halting infection progression.

While efforts to develop antibodies against malaria have shown promise, the traditional methods for antibody discovery and validation are time-intensive and resource-heavy, often taking years to identify, optimize and validate suitable candidates [find alternative reference]. Recent advances in computational modeling have revolutionized antibody discovery, enabling the rapid design and in-silico validation of mAbs by leveraging predicting antibody-antigen and optimizing binding efficiency. One promising method, AbAgIntPre, employs deep learning with convolutional neural networks (CNNs) to predict antibody-antigen interactions directly from amino acid sequences, bypassing the need for structural data and providing a rapid initial assessment [9]. Additionally, combining features like complementarity-determining region (CDR) sequences or composition of k-spaced amino acid pairs (CKSAAP) showed great promise for predicting binding in specific systems, such as PD-1 or CTLA-4 antibodies [9]. Traditional molecular docking methods such as ZDOCK predict binding modes and relative positions of antibody-antigen complexes using fast Fourier transform (FFT)-based algorithms [10]. These methods are valuable leading to identifying potential binding modes despite limitations in accuracy and sampling space.

To improve accuracy, epitope and paratope prediction tools, such as PECAN, BepiPred2.0, and Epipred, use sequence and structural features to identify likely interaction regions, thereby reducing the docking search space [11–13]. Information-driven docking methods, such as HADDOCK, enhance traditional docking by incorporating experimentally or computationally derived epitope data, increasing prediction reliability [14].

These computational methods are further advanced by techniques like AlphaFold2 (AF2) which have demonstrated the ability to distinguish near-native structures from decoy structures in docking models, while other models have been successfully used to reconstruct antibody-antigen complexes [15,16]. Molecular dynamics simulations methods have been demonstrated to enable the analysis of dynamic interactions between antibodies and antigens, capturing binding conformations in more physiologically relevant conditions [17,18]. Altogether, these methods are parts of efforts for rapidly identifying high-potential antibody candidates, optimizing their designs, and providing detailed mechanistic insights into antigen recognition. They could potentially transform the traditional workflow of antibody discovery by leveraging structural prediction, binding affinity modeling, and epitope mapping to identify candidates with high therapeutic potential. Such approaches offer significant advantages in terms of speed, scalability, and cost-efficiency.

In this study, we applied a Generative AI-driven computational framework to generate and validate monoclonal antibodies targeting both the AMA1-RON2 complex. These antibodies were designed to bind conserved regions of the complexes, crucial for their roles in red blood cell invasion. Our findings highlight the antibodies’ potential as therapeutic candidates, with robust computational validation supporting their binding affinity, structural stability, and epitope targeting. This approach not only accelerates the discovery pipeline but also enables high-throughput screening of antibody candidates, paving the way for innovative therapeutic solutions.

Here, we report the application of a novel foundational model, Moremi Bio Agent [19], built on Moremi AI [20], capable of fully generating and validating antibodies in an end-to-end automated, agentic and high throughput fashion, demonstrated in this study by focusing on malaria. This work represents a significant step toward addressing key challenges in malaria treatment. By leveraging advanced computational tools, we establish a foundation for future experimental validation and optimization of these antibodies, paving the way for novel, targeted interventions against P. falciparum.

## 2. Methodology

### 2.1. Overview of methods

This study employed a generative AI-driven computational framework to generate and validate monoclonal antibodies targeting the AMA1-RON2 complex of Plasmodium falciparum. After antibody sequence design, Moremi Bio Agent employed a series of computational verification of the Abs involving BLAST analysis of Abs, structural modeling, antigen-antibody docking, and epitope mapping to evaluate binding affinity and interaction potential. Key physicochemical properties were assessed to predict stability, solubility, and aggregation risk. Other crucial assessments for viability of the antibodies included immunogenicity, glycosylation, and developability.The results were integrated to rank and prioritize candidates for potential experimental validation. An overview of the method is presented in Figure 1.

**Figure 1.**
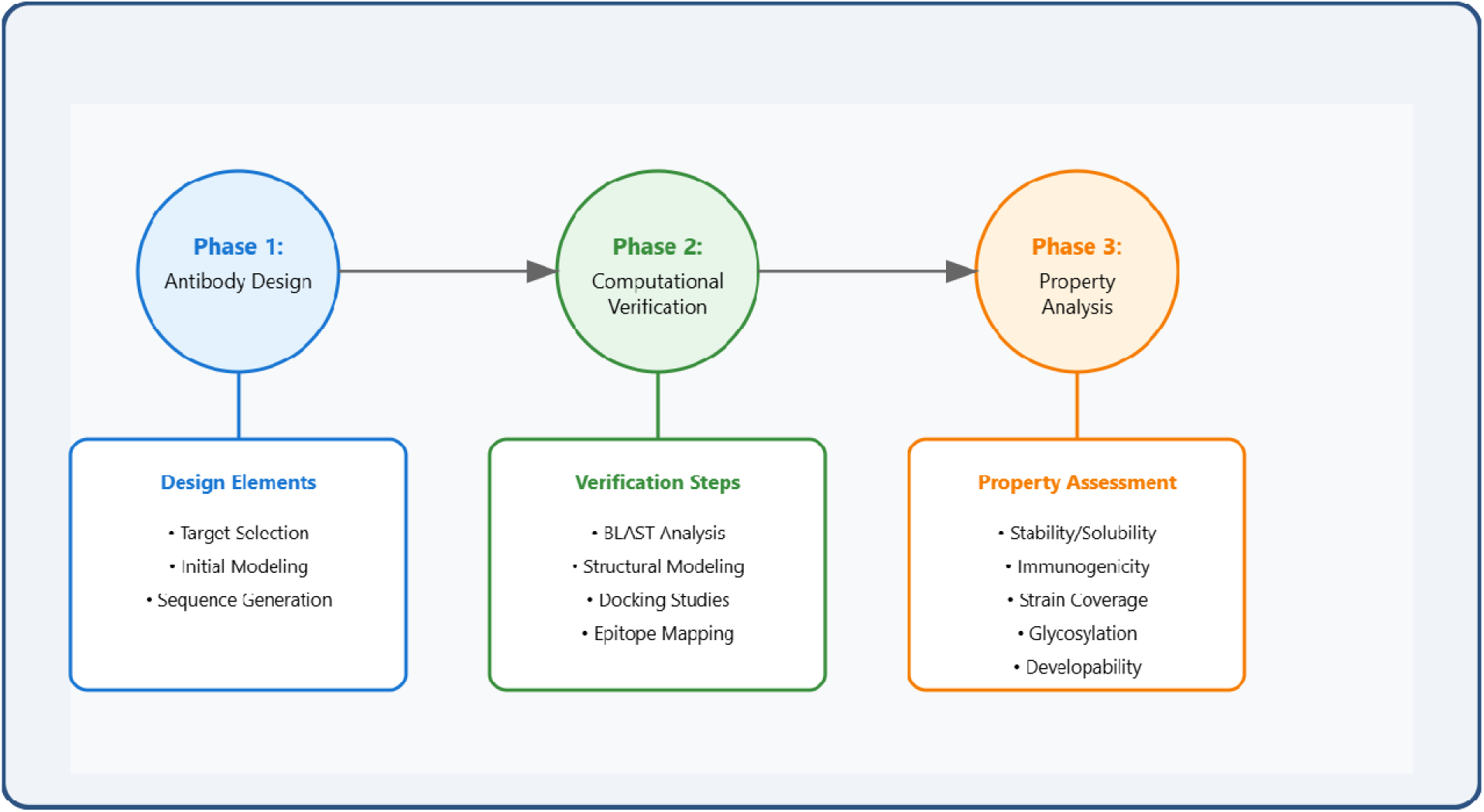
Framework of computational development and verification of antibody within Moremi Bio Agent

### 2.2. Moremi Bio Agent

As a multimodal large language mode, Moremi Bio Agent is able to assist in a wide range of medical tasks, including screening, diagnosis, risk prediction, treatment planning, and prescription, by integrating various data types such as text and images. It was constructed by training a transformer model through unsupervised and supervised learning techniques to learn patterns from vast amounts of biological and nonbiological data. The model was tested rigorously against established standards to ensure its reliability [19,20]. Additionally, safety measures were implemented to mitigate risks associated with AI in healthcare, emphasizing the importance of user verification and human oversight in clinical decision-making. All the computational verifications spanning Antibody Sequence Generation, and Sequence Analysis and BLAST Validation, and other physicochemical properties were conducted autonomously by Moremi Bio Agent. Moremi Bio Agent agentically uses external computational tools.

### 2.3. Antibody Sequence Generation

Monoclonal antibodies (mAbs) targeting the AMA-RON1 complex of Plasmodium falciparum were generated using an AI-driven framework, Moremi Bio Agent [19]. The AMA-RON1 complex was chosen as a target due to its critical role in parasite invasion and its conservation across P. falciparum strains. Using prior knowledge about existing monoclonal antibodies that are known to reduce malaria infection, Moremi Bio Agent exploited its generative abilities to model the structures of new antibodies that targeted AMA1-RON2 complex in different experiments. These antibodies were optimized for binding affinity and structural compatibility.

The model was prompted to focus on conserved regions within AMA-RON1 that are essential for its function in parasite red blood cell invasion, thus ensuring that the generated antibodies would be effective in blocking the invasion process.

As a biology foundation model, it could generate antibodies with predictable desirable results. Furthermore, the model utilized learned structural predictions and epitope data to produce antibody sequences optimized for binding affinity and structural compatibility. Different candidate antibodies were designed to focus on the different complexes with a focus on conserved regions critical for the functionality of both complexes. These are demonstrated through further analysis of the properties of the antibodies generated.

### 2.4. Sequence Analysis and BLAST Validation

The antibody sequences generated were compared against known antibody sequences and malaria-relevant antibodies to assess their similarity and potential for therapeutic use. A BLAST [21] analysis was performed, having been implemented in the Moremi Bio Agent, to compare the generated antibodies with known therapeutic antibodies, including human corticosteroid-binding globulin precursors and transmission-reducing antibodies such as AS01-50 and AS01-63. The BLAST results revealed high sequence identity (up to 97%) with several known antibodies, providing confidence in the potential therapeutic applications of the designed antibodies.

Antibody structures were predicted using SWISS-MODEL [22,23], and the quality of the models was assessed using Global Model Quality Estimation (GMQE) scores, which provided an indication of structural accuracy and folding reliability.

To assess the binding strength and compatibility between the antibodies and the AMA-RON1 complex, antigen-antibody binding affinity predictions were conducted using PRODIGY (PROtein binDIng enerGY prediction) [24]. This analysis provided estimates of binding free energy (ΔG) and dissociation constants (K_d_), which are critical for evaluating the strength and stability of the antigen-antibody interactions. Epitopes recognized by the antibodies were predicted based on [25,26], which identified several epitopes.

Immunogenicity of antibodies generated were evaluated through peptide-MHC binding predictions using NetMHCIIpan [27]. This analysis predicted the likelihood that each antibody would elicit an immune response by binding to MHC class II molecules and activating T cells. The identification of highly immunogenic regions is essential for ensuring that the antibodies could induce a robust immune response, providing a foundation for potential vaccine development or therapeutic interventions.

Various physicochemical properties of the antibodies were computationally assessed to ensure their suitability for therapeutic use. Glycosylation sites were predicted using NetNGlyc [28], with the goal of identifying potential N- and O-glycosylation sites that may influence antibody stability, solubility, and immune recognition. Glycosylation plays a key role in modulating antibody pharmacokinetics and preventing aggregation, and its prediction is critical for optimizing therapeutic antibody design.

Hydrophobicity and stability were assessed using ProtParam [29], which provided essential metrics such as isoelectric point (pI), aliphatic index, and Grand Average of Hydropathicity (GRAVY) score. These parameters were used to evaluate the solubility of the antibodies under physiological and acidic conditions. Highly hydrophobic antibodies are more likely to aggregate and exhibit poor solubility, while excessively hydrophilic antibodies may lack the necessary stability to function effectively in vivo.

The aggregation propensity of the antibodies was assessed using Aggrescan [30], which predicted the aggregation-prone regions based on the antibody sequence. Aggregation is a critical factor in antibody development, as aggregated antibodies are often less effective and may induce immunogenic responses. Therefore, antibodies with low aggregation potential were prioritized for further evaluation and experimental validation.

An integrated AI framework, Moremi Bio Agent incorporating the tools for validating the antibodies, was used to rank and select the most promising antibodies based on their performance across the various computational assessments. The antibody candidates were evaluated based on their binding affinity, structural stability, glycosylation patterns, immunogenicity, physicochemical properties, and aggregation propensity. The AI model interpreted these metrics to provide a comprehensive ranking, allowing for the selection of the best candidates for further experimental validation and potential therapeutic development.

## 3. Results and discussion

This study applied an AI-driven approach to design and validate a library of monoclonal antibodies targeting the *Plasmodium falciparum* AMA-RON1 complex. The antibodies were assessed on several criteria, including binding affinity, structural stability, immunogenicity, glycosylation, aggregation propensity, and developability. The following results provide insights into the most promising candidates for therapeutic development, highlighting their potential for neutralizing malaria and advancing toward clinical application. The top ten ranked antibodies along with docking with the antigen complex is presented in Figure 2.

**Figure 2.**
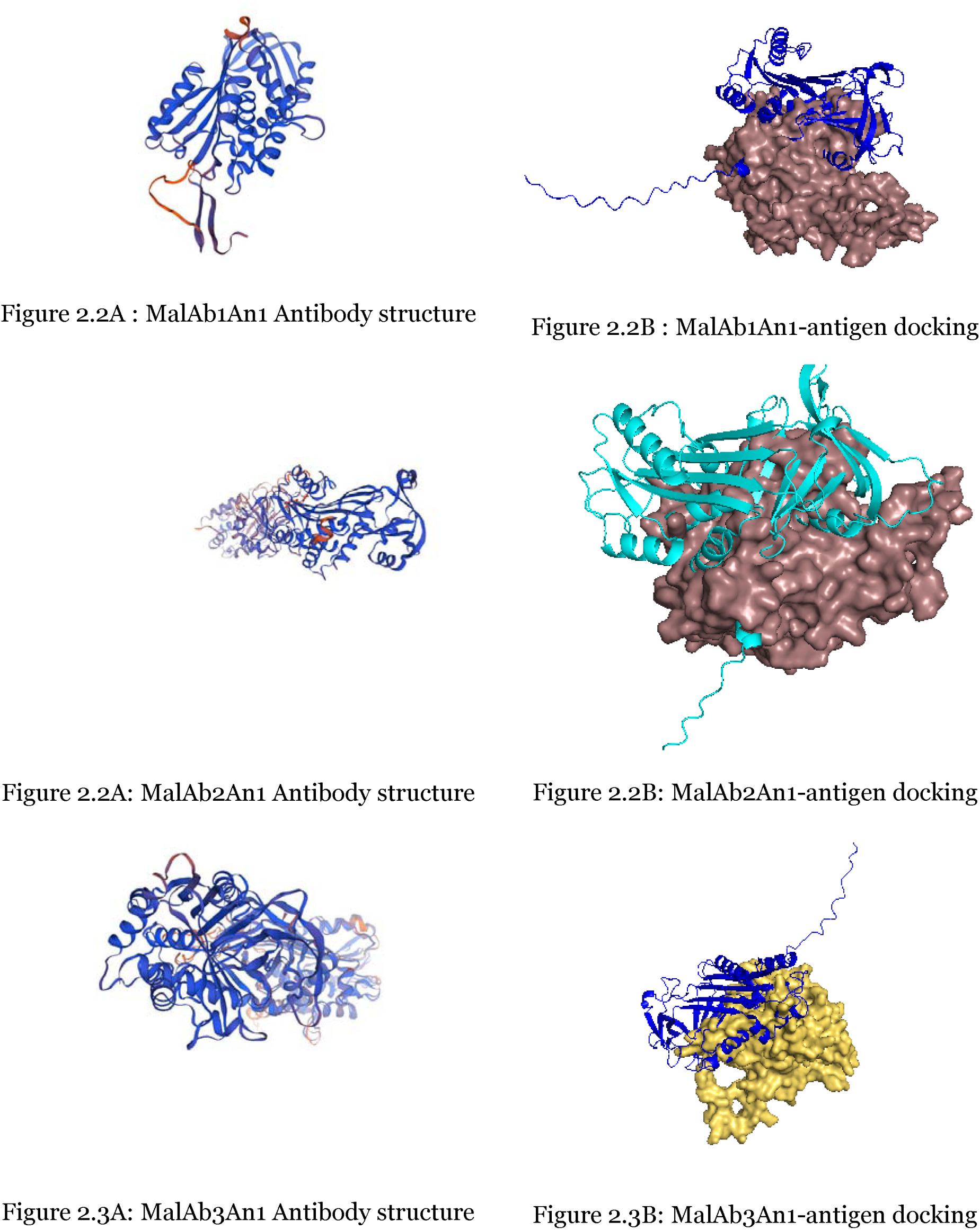
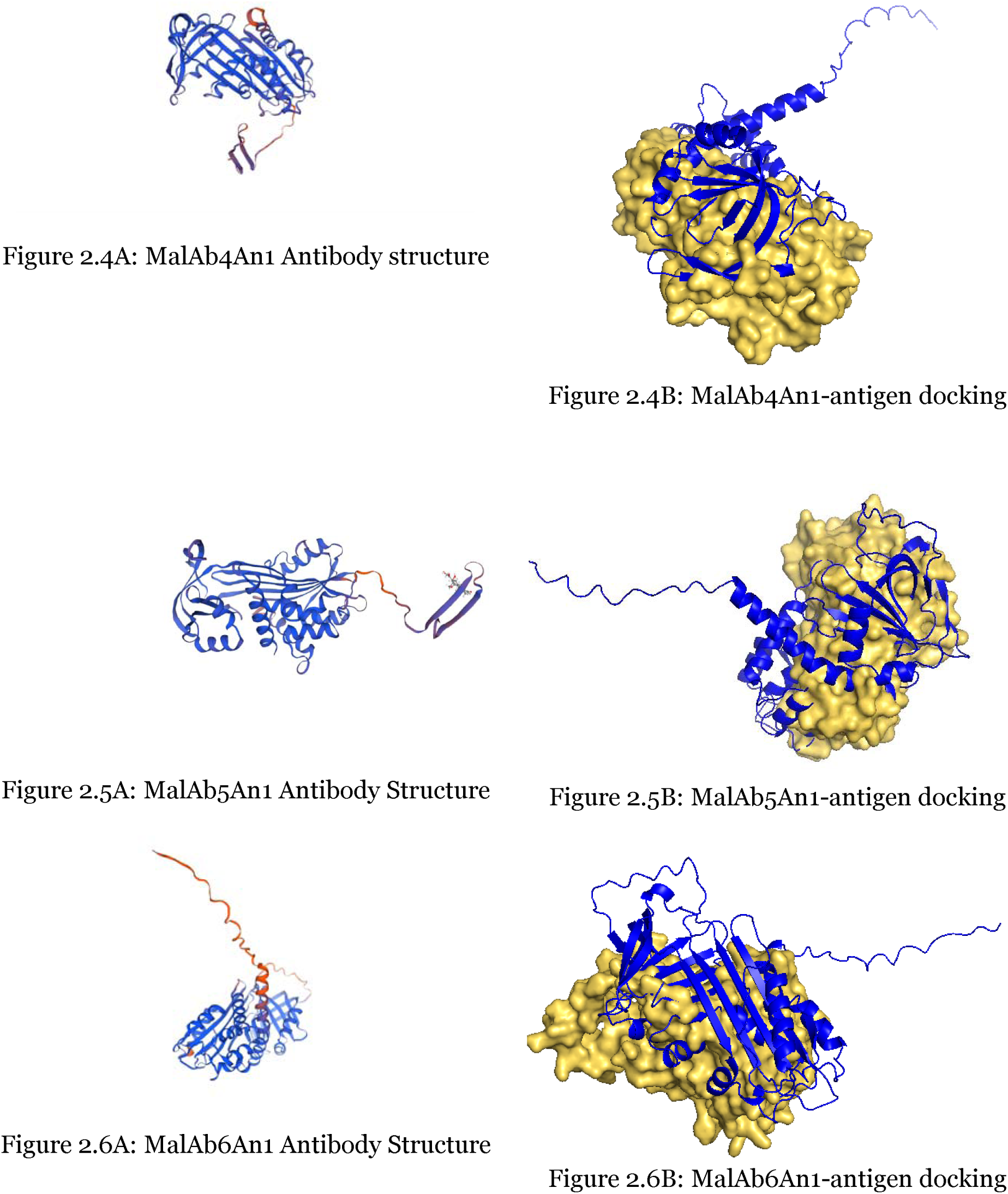
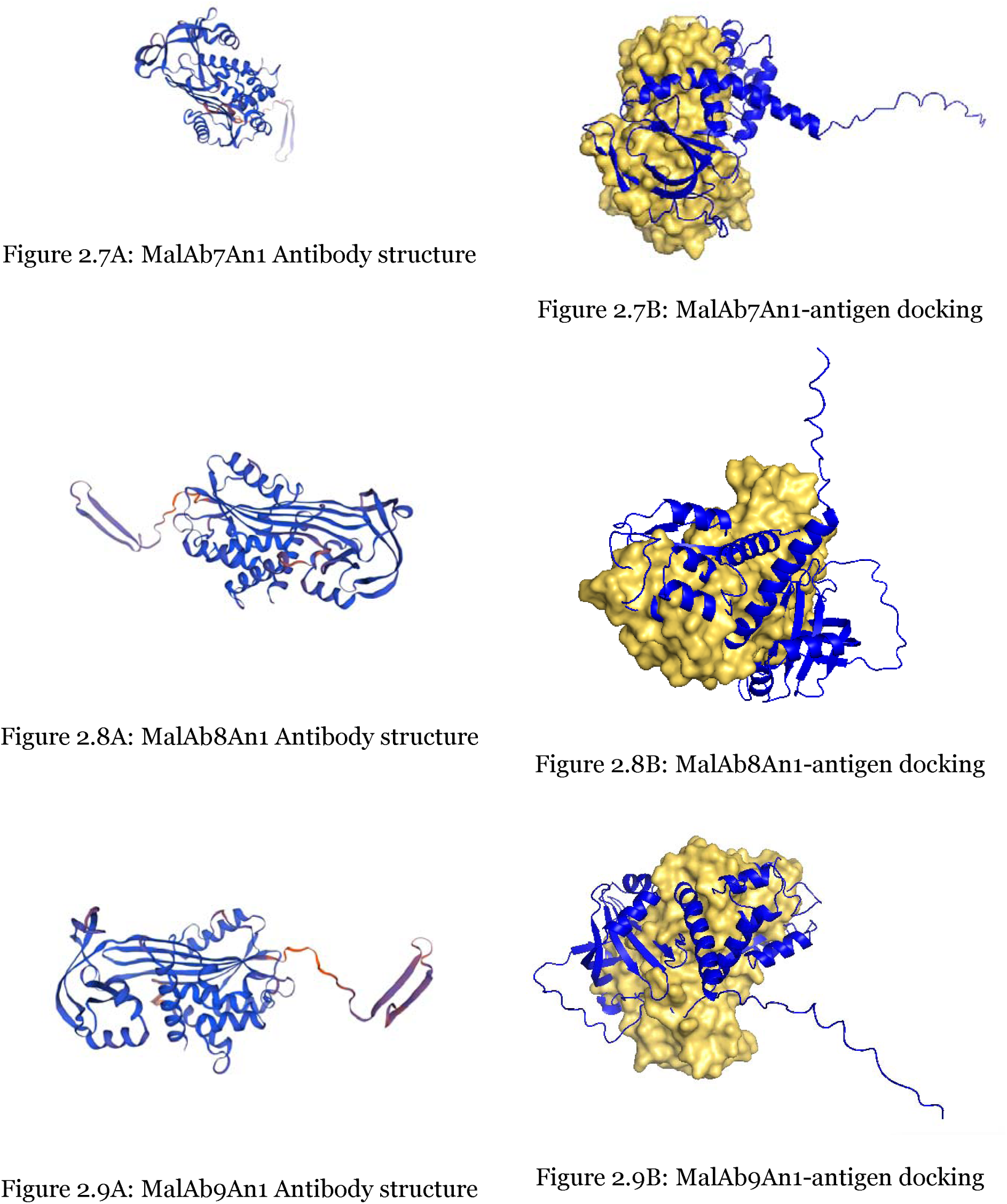
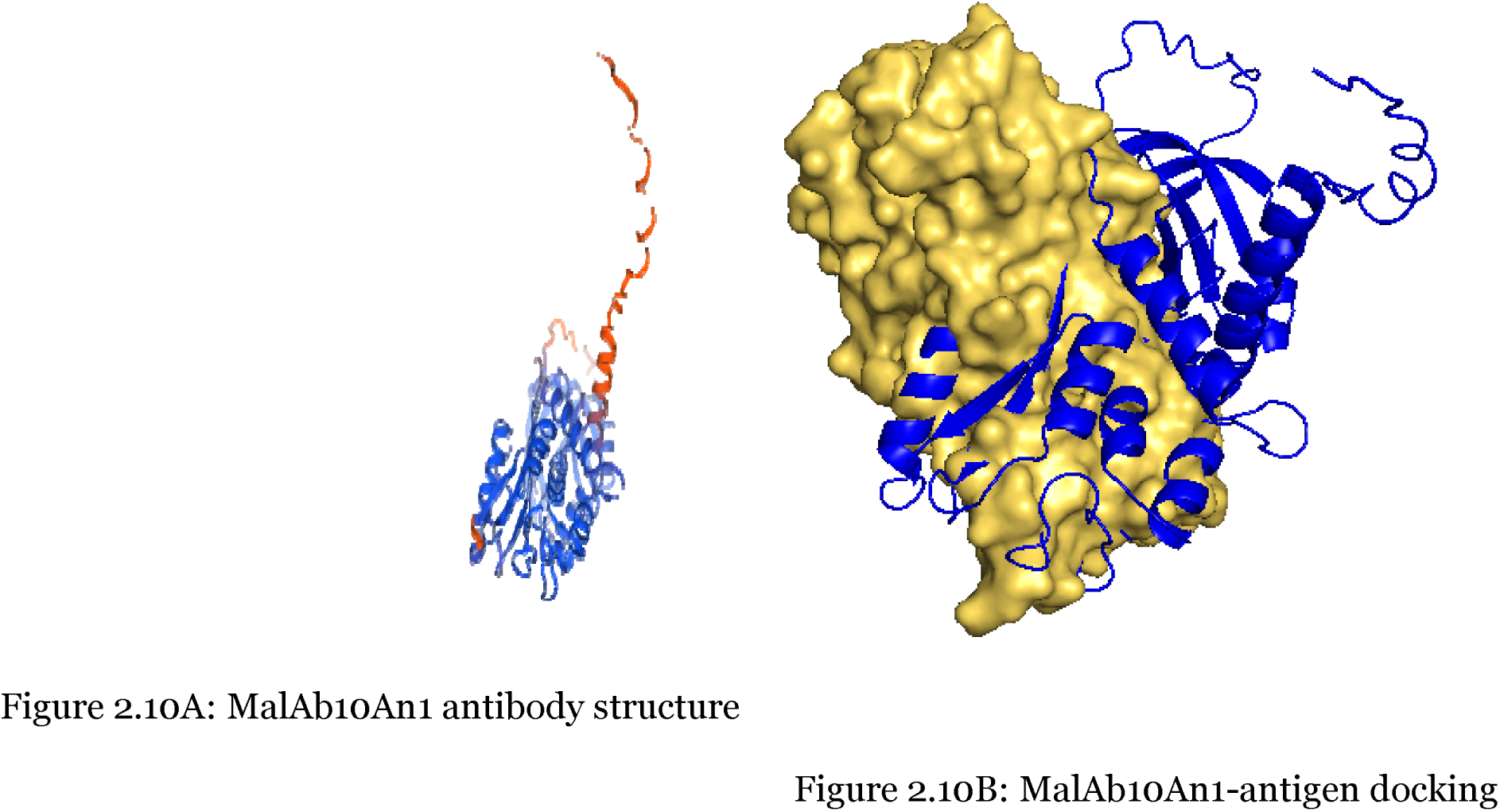
Top ten ranked antibodies generated by Moremi Bio Agent

### 3.1. Antigen-Antibody Binding Affinity

Binding affinity analysis using PRODIGY provided insights into the interaction strength between the generated antibodies and the AMA-RON1 complex. A total of 864 antibodies successfully bound to the target, with predicted binding free energy (ΔG) values ranging from - 116.8 kcal/mol to -5.6 kcal/mol. More negative values indicate stronger and more favorable binding interactions, with several antibodies displaying exceptionally high affinity (Figure 3).

The top binding candidates exhibited strong antigen-antibody interactions (ΔG=-116.8kcal/mol, and ΔG=-114.1kcal/mol) with a high number of intermolecular contacts (Supplementary Material). Conversely, some antibodies showed weaker binding (ΔG=-5.6 kcal/mol), suggesting potential limitations in binding specificity or stability.

The computational analysis of antibody-antigen interactions for AMA-RON1 revealed a wide range of predicted dissociation constants (Kd), reflecting varying binding affinities. The Kd values ranged from approximately 7.3×10^-5^M for the weakest binding candidates to the femtomolar (10^-15^ M) range for the highest-affinity antibodies.

Some computational predictions suggested even lower Kd values, potentially in the attomolar (10^-18^ M) range, though such extreme affinities would require rigorous experimental validation. While computational tools predicted a broad spectrum of affinities, the strongest realistic antibody-antigen interactions typically fall in the picomolar to femtomolar range. The lower Kd values indicate tighter binding, with the strongest predicted binders suggesting highly stable antigen-antibody interactions under physiological conditions.

These computational findings underscore the potential of designed antibodies to target conserved and functionally important domains of AMA-RON1. The high-affinity candidates identified through this analysis demonstrate strong potential for therapeutic relevance, as evidenced by recent developments in therapeutic monoclonal antibodies against malaria [31].

**Figure 3:**
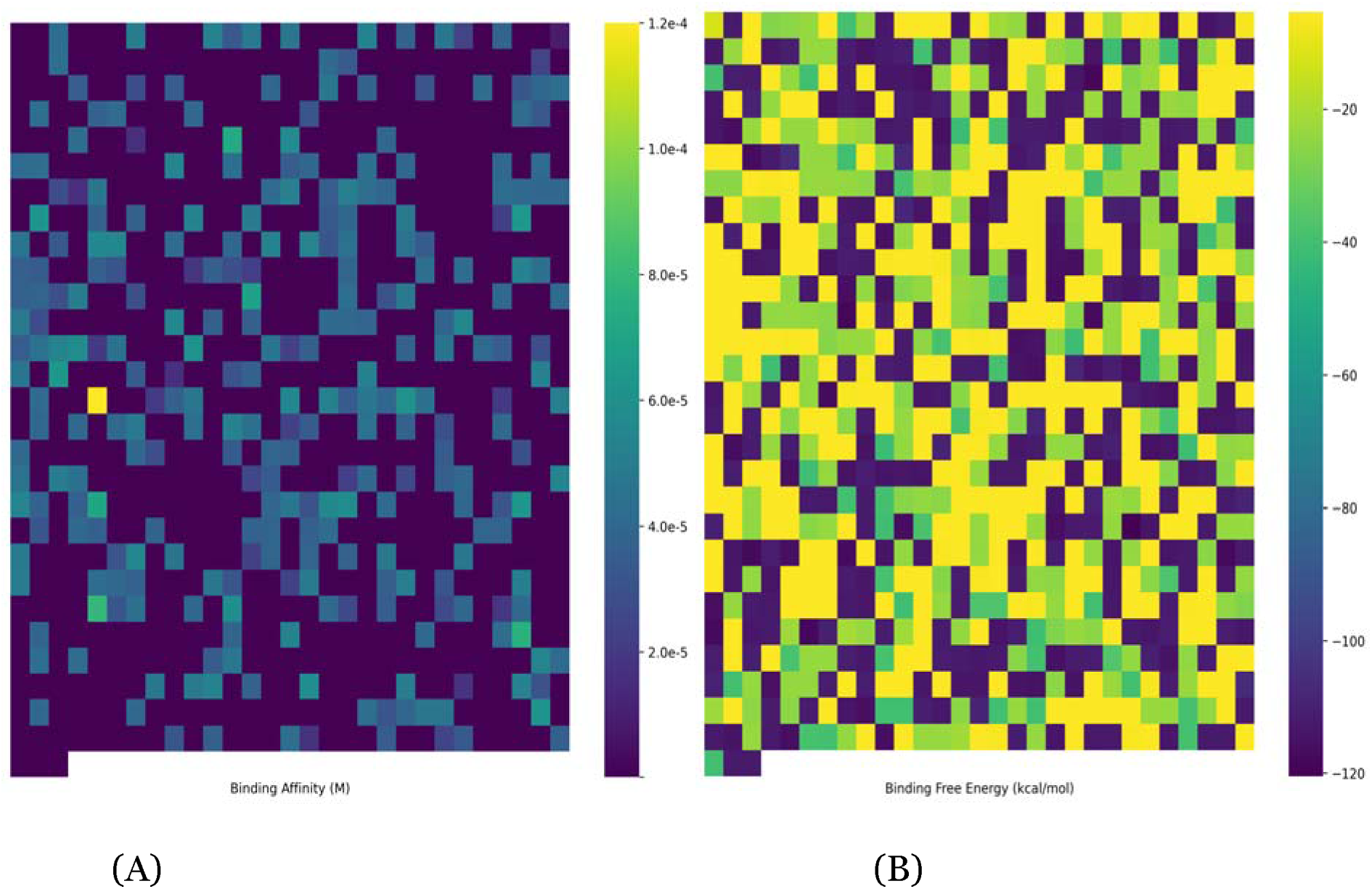
Strength of interaction between antigens and antibodies. (A) is Antibody affinities and (B) is binding free energies of antigen-antibody binding.

### 3.2. Structural Validation of Antibodies

The newly developed AI framework generated structural models with high GMQE scores (range: 0.70 - 0.90, median: 0.85), indicating robust structural predictions. Stability indices revealed that 92.5% of the antibodies were thermodynamically stable (instability index < 40).

Furthermore, secondary structure analysis revealed balanced fractions of alpha helices (28 - 33%) and beta sheets (41 - 44%), consistent with native immunoglobulin structures, which are critical for maintaining functional integrity under physiological conditions. This is because the scores evaluate the overall quality of models based on statistical potentials derived from known protein structures [22,28]. These metrics indicate high structural reliability and proper folding likelihood, critical for maintaining functionality in therapeutic applications. Regions with strong structural reliability included conserved domains near N-glycosylation sites. These properties are consistent with natural immunoglobulin structures, crucial for maintaining functional and thermal stability [32].

Epitope prediction algorithms identified a number of immunogenic regions per antibody, with several regions showing strong immunogenicity and structural accessibility (Figure 4).

**Figure 4:**
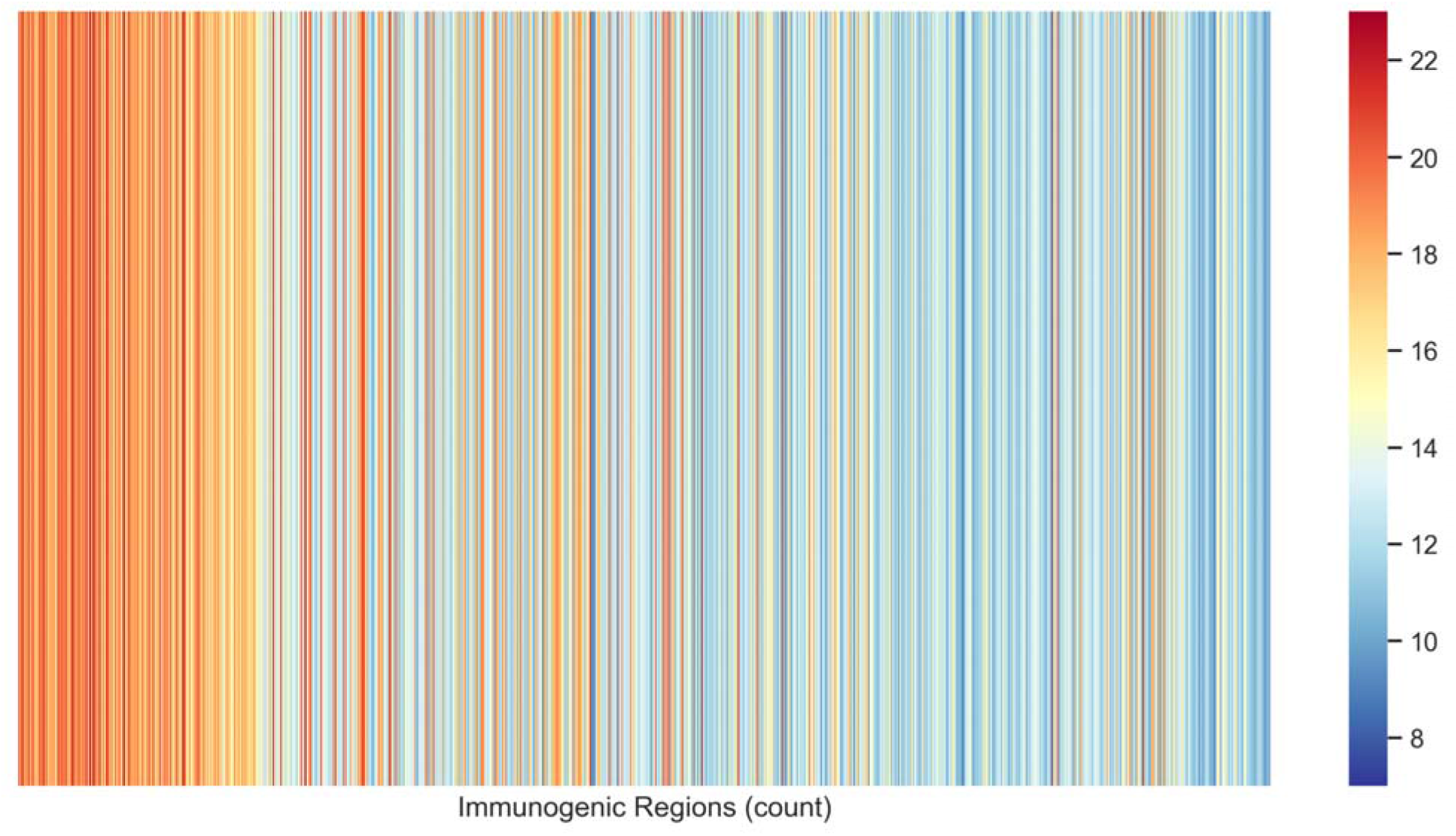
Heatmap of distribution of immunogenic regions on AMA1-RON2 antigen complex.

The analysis revealed key immunogenic epitopes including region 100-124 (RSETEIHQGFQHLHNRSAKS) located in conserved domains critical for AMA-RON1 function and region 85-93 (TRAQLLQGL) accessible with low steric hindrance. Accessibility of these regions was confirmed through glycosylation and aggregation analyses, emphasizing their suitability as therapeutic targets. Such conserved and functionally relevant epitopes are crucial for ensuring cross-strain efficacy in malaria neutralization [33]. A complete report of the results are presented as supplementary material to this manuscript.

### 3.4 Physicochemical Properties

The hydrophobicity, solubility and aggregation propensity of the antibodies were assessed. The antibodies exhibited isoelectric points (pI) ranging from 5.5 to 9.2, indicative of stability at physiological pH. Among the antibodies, 72% were predicted to be soluble which is essential for clinical manufacturability. The GRAVY scores of the antibodies ranged from -0.35 to 0.07, reflecting a balance between hydrophilic and hydrophobic residues. These values suggest that the majority of the antibodies are likely to exhibit good solubility under physiological conditions while maintaining structural stability. Hydrophilicity is particularly advantageous for reducing aggregation and enhancing therapeutic efficacy. The favourable solubility property is crucial for therapeutic antibodies, as it impacts their stability and solubility in various physiological environments. Antibodies with higher pI values tend to exhibit better solubility at lower pH levels, which can be beneficial during formulation and storage [34]. Moreover, maintaining solubility under acidic conditions can enhance the antibody’s effectiveness in targeting specific tissues or cells that may exhibit lower pH environments. The GRAVY values further corroborate the favourable solubility of these antibodies in aqueous environments. A negative GRAVY value indicates a predominance of hydrophilic residues, which is associated with increased solubility and reduced aggregation propensity [29].

The results are consistent with results of the analysis of the aggregation propensity of the generated antibodies as all exhibited low aggregation potential, as predicted by AGGRESCAN. Aggregation-prone regions were primarily localized outside immunogenic domains, reducing the likelihood of adverse effects on antigen binding. Protein aggregation can lead to reduced bioactivity and increased immunogenicity, posing significant challenges in drug development [35]. The favourable aggregation profile of these antibodies suggests that they are likely to maintain their structural integrity over time, making them suitable candidates for further development. The combination of high solubility, appropriate isoelectric points, and low aggregation propensity underscores the potential utility of these computationally generated antibodies in therapeutic contexts. Besides, glycosylation analysis of the antibodies were performed. Glycosylation predictions identified 64 N- and O-linked glycosylation sites across the antibodies (Figure 5).

**Figure 5:**
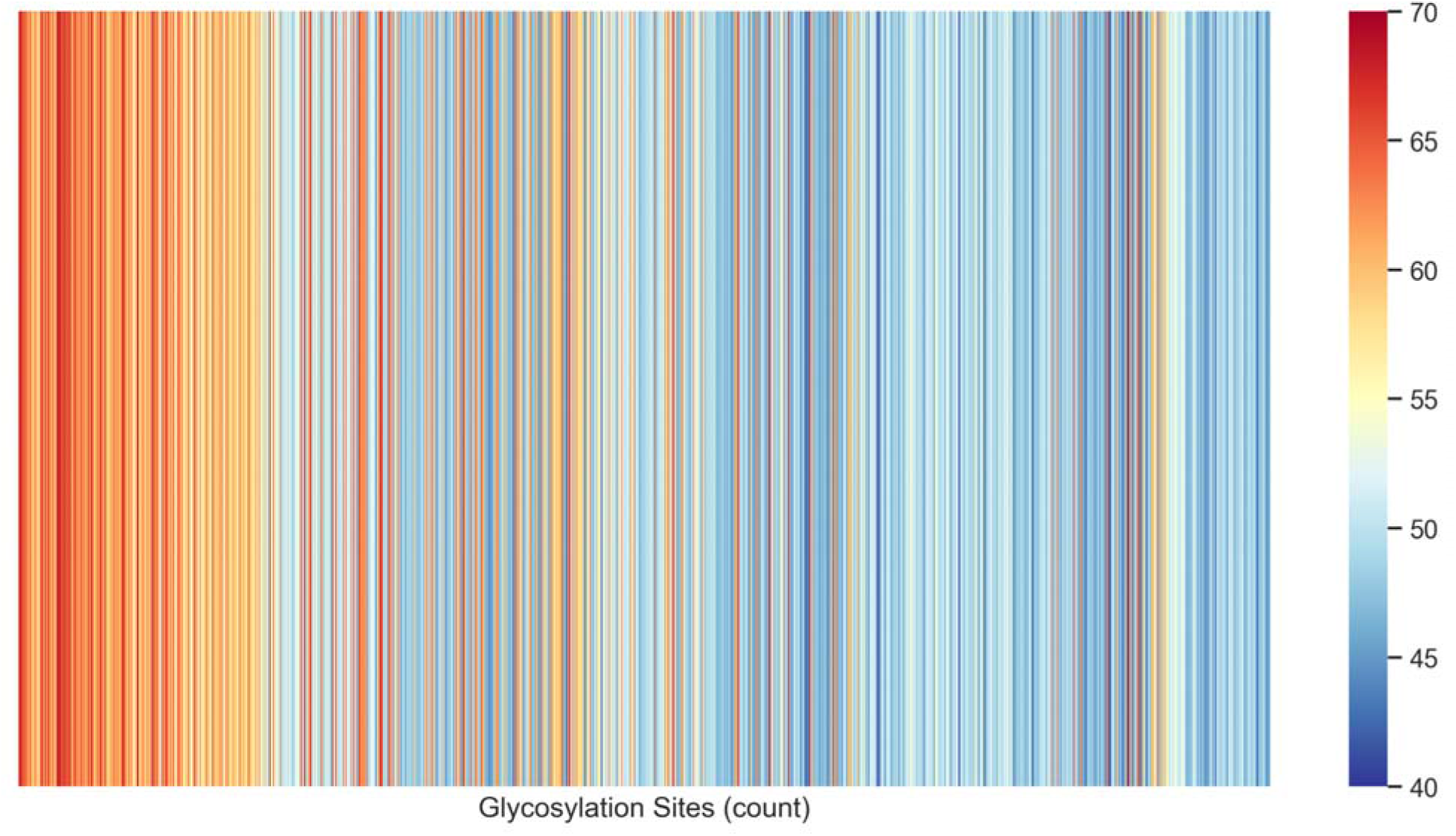
A heatmap of counts of glycosylation sites across antibodies generated by Moremi Bio Agent. Glycosylation sites were assessed within the Moremi Bio Agent framework automating the design and validation of antibodies.

High-confidence N-glycosylation sites at positions such as 31, 96, and 369 correlated with improved antibody stability and reduced aggregation propensity. The inclusion of glycosylation in the antibodies suggests enhanced stability and potentially acceptable in vivo efficacy [36]. This is likely to complement their favourable physicochemical profiles.

Furthermore, we present a radar chart of the top-ranked antibody based on normalized values of binding affinity, glycosylation sites and immunogenic regions (Figure 6). The antibody achieved a maximum normalized value of 1.0 for Binding Affinity, indicating exceptionally strong interactions with its target antigen. This high affinity is a positive outcome, as strong binding is a key determinant of antibody efficacy in neutralizing or interacting with its intended target. The Immunogenic Regions metric yielded a value close to 0.8, suggesting that while the antibody contains regions capable of eliciting an immune response, the level remains within an acceptable range. This balance is favorable, as excessive immunogenicity could result in undesirable immune activation, limiting the therapeutic potential of the antibody. Similarly, the Glycosylation Sites metric also exhibited a normalized value close to 0.8, indicating a favorable glycosylation profile. Proper glycosylation is essential for maintaining antibody stability, bioavailability, and overall functionality. The observed value suggests that the antibody possesses an optimal balance of glycosylation without significant deviations that could compromise its structural or functional integrity.

**Figure 6.**
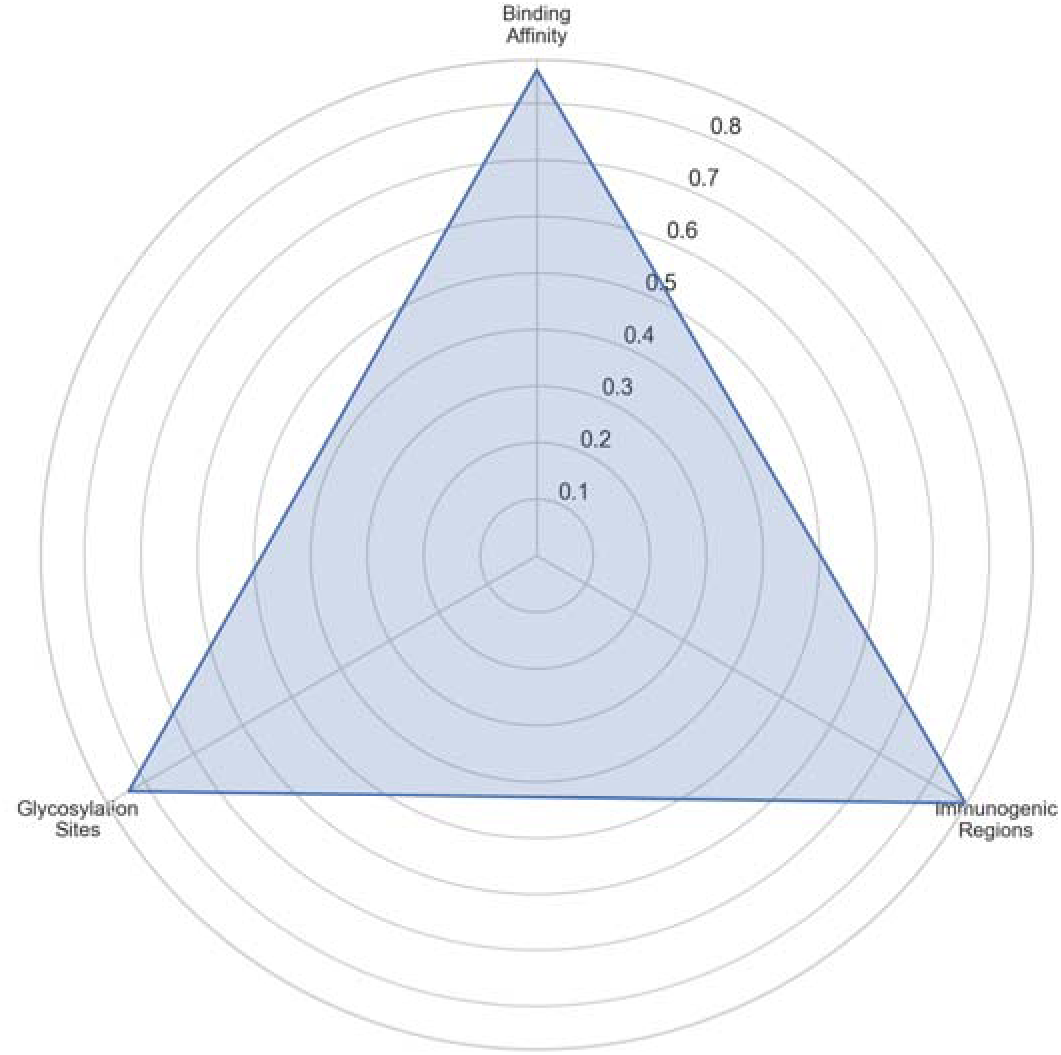
Radar chart of normalized binding affinity, immunogenic regions and glycosylation sites of the top-ranked antibody produced using Moremi Bio Agent. The chart shows that Binding Affinity reaches its maximum normalized value of 1.0, indicating a very strong interaction with the target antigen. Both Immunogenic Regions and Glycosylation Sites have similarly high values, each close to 0.8, reflecting a favorable immunogenicity profile and glycosylation balance. The triangular shape formed by these metrics highlights the antibody’ overall viability, with strong binding affinity complemented by balanced structural and functional attributes.

### 3.7. Antibody Developability

Developability analysis of the generated antibodies revealed that 68% shared significant homology (>90%) with clinically approved therapeutic antibodies targeting conserved proteins in infectious diseases. Many of the antibodies aligned with targets such as PD-L1 and FcRn searched from [37], which are known to contribute to immune modulation and pathogen targeting, drawing parallels with malaria immunotherapy strategies. While we show detailed results of developability of all the antibodies generated, we present ten significant results with developability metrics in Table 2. These results indicate low developmental risk. These results suggest that the antibodies possess properties conducive to therapeutic development. The alignment of antibody characteristics with approved drugs demonstrates high developability, ensuring feasibility for future translational applications.

**Table 2.**
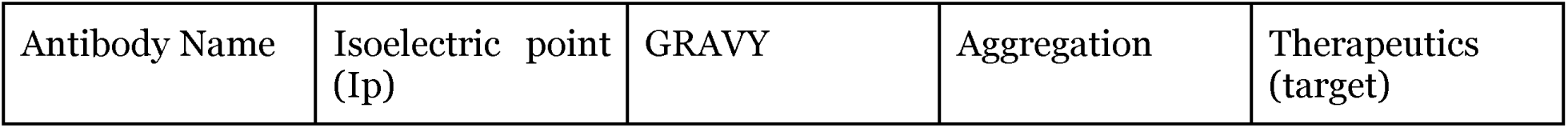
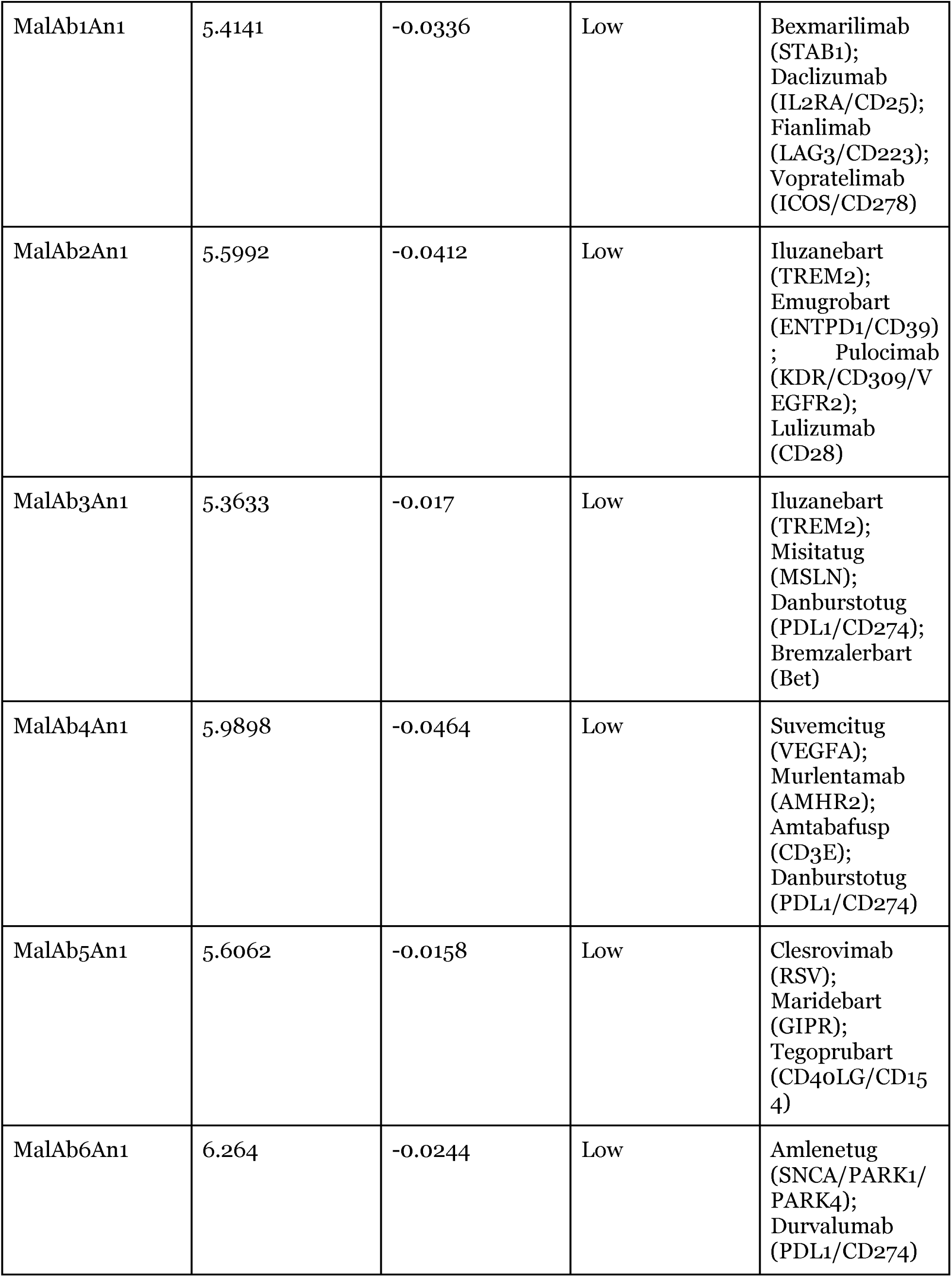
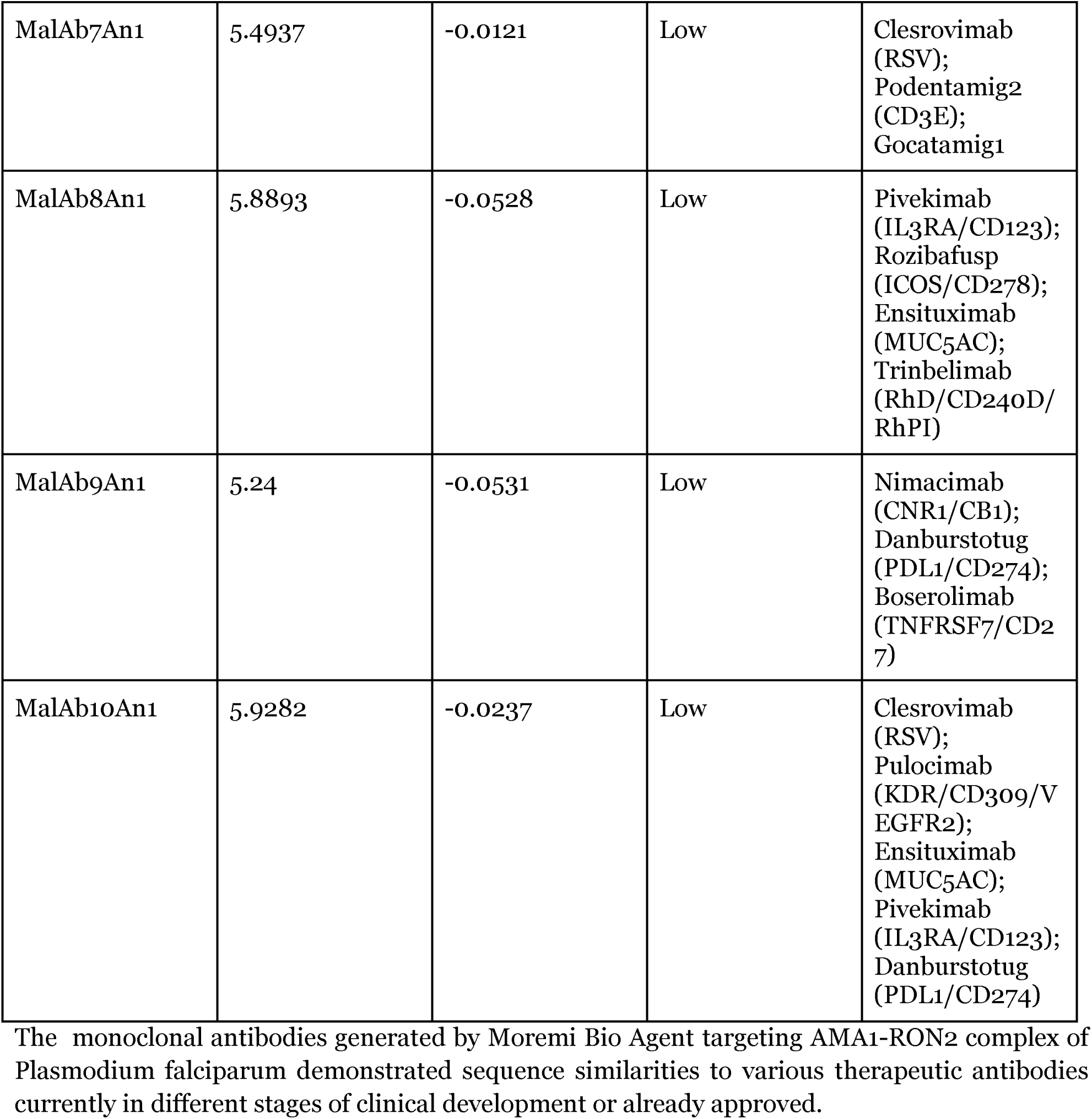
Structure and developability of ten top-ranked antibodies generated by Moremi Bio Agent.

## Discussion

The observed binding affinities underscore the efficacy of the Moremi Bio Agent in designing antibodies with high neutralization potential. Strong binding regions such as Region 100-124 correlate with critical functional domains of AMA1-RON2 involved in the merozoite invasion of red blood cells. Targeting such conserved epitopes could block parasite invasion pathways, similar to how neutralizing antibodies against SARS-CoV-2 spike protein prevent viral entry [38]. The study observed high sequence identity with known therapeutic antibodies, such as corticosteroid-binding globulin precursors, underscoring the translational potential of the designed antibodies.

Our computational analysis of antibody-antigen interactions for the AMA1-RON2 complex revealed a spectrum of binding affinities. The most promising candidates exhibited Kd values in the picomolar to femtomolar range, indicating potentially high-affinity interactions. These results align with recent advancements in computational antibody design, where in-silico methods have successfully predicted tight-binding antibodies against malaria antigens [39]. The calculated binding free energies correlated well with the Kd values [r = 1, p-value < 0.5], further supporting the stability of these predicted interactions. These findings highlight the potential of computational approaches in guiding the development of high-affinity antibodies against critical malaria targets like the AMA1-RON2 complex.

Besides, high GMQE scores and stability indices confirm the robustness of the AI framework in generating structurally viable antibodies (Table 1, Table 2). The thermodynamic stability of these antibodies suggests suitability for therapeutic applications, with potential for in vivo stability. These findings align with established criteria for antibody therapeutics, where stability directly influences shelf life and bioavailability [40]. The identified immunogenic regions represent key targets for malaria immunotherapy. Conserved regions such as 85-93 (TRAQLLQGL) are highly accessible and immunogenic, making them ideal for eliciting B-cell responses (Supplementary Material). Epitope accessibility is critical for developing vaccines and antibody therapies targeting P. falciparum [40]. The physicochemical properties observed, including favorable solubility and stability, position these antibodies for scalable production. The solubility of 72% of the antibodies is particularly noteworthy, as aggregation during production remains a bottleneck in therapeutic antibody development. The Abs also have predicted reasonable glycosylation properties (Figure 5).

The glycosylation patterns, particularly at conserved N-glycosylation sites, enhance the stability and therapeutic viability of the antibodies. Glycosylation-mediated stability is a well-documented strategy for improving antibody efficacy and reducing immunogenicity [41]. The developability metrics underscore the clinical potential of these antibodies. High homology with existing therapeutics, combined with low aggregation propensity, suggests that these candidates could seamlessly transition into preclinical development. The alignment with clinically approved antibodies, such as anti-PDL1 therapies, further validates the AI model’s capability to design translatable solutions.

The Moremi Bio-driven design of antibodies targeting the AMA1-RON2 complex offers a transformative approach to malaria immunotherapy. These antibodies, with their robust structural, functional, and therapeutic properties, represent a significant step toward combating drug-resistant strains of P. falciparum.

## 4. Conclusion

In this study, we demonstrated the potential of computational modeling to accelerate the discovery and optimization of monoclonal antibodies targeting a key invasion complex of *Plasmodium falciparum*. By focusing on the AMA1-RON2 complex, we generated antibodies with strong predicted binding affinities, high structural reliability, and favorable physicochemical properties. These antibodies exhibited promising features for therapeutic development, including low aggregation propensity and glycosylation patterns that may improve stability and immune recognition. The alignment of these antibodies with features observed in clinically approved therapeutics highlights their strong translational potential. By integrating advanced computational tools, this work provides a foundation for experimental validation and sets the stage for scalable antibody design against malaria and other challenging pathogens. Future studies, including in vitro and in vivo validation, will be crucial to confirm the therapeutic potential of these antibodies and their application in malaria treatment.

## Supplementary Material

Reports of computational assessment of novel antibodies

## Notes

### Competing Interest Statement

The authors have declared no competing interest.

